# Linking phenology, harvest index and genetics to improve chickpea grain yield

**DOI:** 10.1101/2024.04.23.590839

**Authors:** R. Gimenez, L. Lake, M. C. Cossani, R. Ortega Martinez, J. E. Hayes, M. F. Dreccer, R. French, J. L. Weller, V. O. Sadras

## Abstract

Phenology is critical to crop adaptation. We grew 24 chickpea genotypes in 12 environments to analyse: the environmental and genotypic drivers of phenology; associations between phenology and yield; and phenotypes associated with allelic variants of three flowering related candidate loci: *CaELF3a*; a cluster of three *FT* genes on chromosome 3; and a region on chromosome 4 with an orthologue of the floral promoter *GIGANTEA*. A simple model with 3 genotype-specific parameters explained the differences in flowering response to daylength. Environmental factors causing flower abortion, such as low temperature and radiation and high humidity, led to a longer flowering-to-podding interval. Late podding associated with poor partition to grain, limiting yield in favourable environments. Sonali, carrying the early allele of *Caelf3a* (*elf3a*), was generally the earliest to set pod, had low biomass but the highest harvest index. Genotypes combining the early variants of *GIGANTEA* and *FT* orthologues *FTdel*, where a deletion in the intergenic region of *FTa1-FTa2* was associated with slow development, usually featured early reproduction and high harvest index, returning high yield in favourable environments. We emphasise the importance of pod set, rather than flowering, as a target for breeding, agronomic, and modelling applications.

**Highlight:** This paper analyses the environmental and genetic controls of chickpea phenology and its effects on grain yield, in a multi-environment trial including 24 genotypes with varying combinations of flowering related genes.

## Introduction

Chickpea (*Cicer arietinum* L) is a globally important pulse, valued as an affordable and nutrient-dense source of plant-based protein (Jukanti *et al*., 2012; Singh *et al*., 2021). It ranks second in annual production among pulses, surpassed only by dry beans, with 18.1 M t in 2022 (Anwar *et al*., 2022; FAOSTAT, 2024; Singh *et al*., 2021). Originating in the Fertile Crescent around 7,500 years ago, chickpea was initially cultivated as a winter crop in Mediterranean climates with mild, wet winter and hot, dry summer (Redden and Berger, 2007). However, early in its cultivation, it shifted from an autumn-sown to a spring-sown crop, presumably to escape the devastating Ascochyta blight disease (Abbo *et al*., 2003b). This shift involved significant alterations in key adaptation traits, including a putatively reduced vernalisation response, reduced cold tolerance, and increased reliance on stored soil moisture, shaping its subsequent habitat range (Abbo *et al*., 2003a; Berger *et al*., 2023; Berger, 2007).

Contemporary chickpea is adapted to a wide range of environments including semi-arid regions with low rainfall and high temperature (*i.e.,* Africa, South-East Asia, Middle East) and subtropical and temperate regions with warm, wet summer and mild, dry winter (*i.e.,* northern Australia, South America) (Malhotra and Singh, 1991). In recent decades, its global relevance has grown due to the combination of a rising demand for plant-based food, favourable international prices, and recognised benefits for crop rotations. These factors made chickpea an important component not only for subsistence systems but also for extensive agricultural systems of countries like Australia, USA, and Canada (Berrada *et al*., 2007; Merga *et al*., 2019). There are two main commercial types of chickpeas: ‘kabuli’, which evolved in the Middle East, and is mainly cultivated as a summer crop in the Mediterranean area, the Near East, and central and south America; and the smaller-seeded ‘desi’ types from India, which are mostly grown as a winter crop in the semi-arid tropics, sub-tropics of East Africa and Oceania (Ellis *et al*., 1994; Singh *et al*., 2021).

In Australia, chickpea production was incipient in the early 1980s and reached over 1 M t in 2022 (FAOSTAT, 2024). Currently, Australia is the largest global exporter of desi chickpea (Anwar *et al*., 2022). The crop is grown mainly in the warm temperate north-east region of southern Queensland and northern New South Wales (Whish *et al*., 2007), but has expanded to Mediterranean-type environments of Western and South Australia where drier and cooler conditions constrain crop yield (Lake *et al*., 2016; Richards *et al*., 2020). Despite its growing importance and research and breeding efforts, chickpea yield gains have been limited (Richards *et al*., 2022), and grower yields remain low and highly variable (Anwar *et al*., 2022). Estimated yield gaps (*i.e.* the difference between farmer yield and water-limited yield potential) are in the order of 50 -75%, highlighting the potential for agronomic practices to close these gaps (Mawalagedera and Brand, 2022; Peake *et al*., 2021).

The crop-specific critical period is a relatively short phase during the crop cycle when stress is more likely to reduce grain number and crop yield (Carrera; *et al*., 2023). In chickpea, the reported critical period for yield determination spans from 300 °Cd before to 500 °Cd after flowering, peaking at the onset of pod set (Lake and Sadras, 2014). Crop growth rate during this period has been linearly related to crop yield (Lake and Sadras, 2016) whereas the exposure to severe stress can cause yield losses as high as 80% (Lake and Sadras, 2014). Therefore, pairing cultivar and sowing time to ensure that the critical period coincides with the best growing conditions is critical to yield. This requires understanding of both the phenology of available cultivars and the characteristics of the growing environments (Singh *et al*., 2021).

Phenology is more complex and less understood in chickpea than in other pulses and cereals (Singh *et al*., 2021). In addition to temperature and daylength (Roberts *et al*., 1985), soil water availability also affects chickpea phenology (Chauhan *et al*., 2019; Li *et al*., 2022; Singh and Virmani, 1996). Warm temperatures, long days and water deficit hasten chickpea’s rate of development, with interactions among these environmental factors, and genotype-dependent variation in the response (Daba *et al*., 2016; Richards *et al*., 2020; Siddique *et al*., 2000). Factors like chilling temperature, low radiation or high humidity can cause abortion of flowers and developing pods, further delaying the onset of flowering and/or pod setting (Berger *et al*., 2012; Clarke and Siddique, 2004; Croser *et al*., 2003; Jettner *et al*., 1999; Srinivasan *et al*., 1998; Verghis, 1996). A better understanding of how environmental factors interact to regulate the phenology of chickpea cultivars and how the resulting growing conditions during the critical period affect grain yield, would support improved breeding strategies and crop management tailored for specific environments or climate scenarios (Whish *et al*., 2007).

The combination of classical mapping and comparative genomic studies has elucidated several genes associated with flowering time in chickpea. These include the major floral repressor *CaELF3a* on chromosome 5, a cluster of three *FT* genes on chromosome 3, and a region on chromosome 4 with an orthologue of the floral promoter *GIGANTEA* (Atieno *et al*., 2021; Ortega *et al*., 2019; Ridge *et al*., 2017). Besides these, three major loci and multiple QTL without suggested candidates have been reported (Gaur *et al*., 2015; Mallikarjuna *et al*., 2017; Varshney *et al*., 2014). The allelic characterisation of chickpea at candidate genes may be useful in breeding.

We grew 24 chickpea genotypes in diverse Australian environments spanning winter-rainfall regimes in the south-east and west, and summer-rainfall areas in the northeast (Lake *et al*., 2016). Our aim was to analyse: i) the environmental and genotypic drivers of their phenology; ii) associations between phenology and yield and its components; and iii) associations between agronomically relevant field phenotypes and the allelic expression of the three main candidate genes outlined above.

## Materials and Methods

### Experimental design and crop husbandry

Field experiments were conducted in 12 environments consisting of three sites (Kapunda, -34.40, 138.86, South Australia; Merredin, -31.49, 118.23, Western Australia, and Gatton, -27.57, 152.36, Queensland), 2 seasons (2021 and 2022) and 2 times of sowing. Crops were sown on 5 May and 29 June 2021, and 10 May and 15 July 2022 at Kapunda; on 19 May and 25 June 2021, and 27 May and 23 June 2022 at Merredin; and on 24 May and 22 June 2021, and 16 June and 20 July 2022 on Gatton. Experiments included 24 genotypes, 19 desi and 5 kabuli, including mostly commercial varieties and 2 advanced breeding lines (Table 1), that were allocated to a randomised complete block design with three replicates. Plots size and configuration varied across sites to account for local practice (6 rows, 0.22-0.25 m apart at Kapunda and Merredin; 4 rows, 0.35 m apart at Gatton), with a common target of 50 pl m^-2^ for desi and 30 pl m^-2^ for kabuli. Seeds were pre-treated with P – Pickel T fungicide to minimize the risk of *Ascochyta* blight and inoculated with Group N rhizobia at sowing. Best local practices were used to manage weeds, pests, and diseases.

**Table 1:** Chickpea genotypes detailing their type, label (between brackets), allelic composition for *ELF3a*, *GI* and *FTdel* (where asterisks indicate the early allele); allelic group based on the presence of one or more early alleles, and the parameters of their daylength response. Daylength response parameters are: DLc: the threshold daylength below which flowering time responds to daylength; DLs, daylength sensitivity, quantified as the rate of change in flowering time per unit change in daylength above the threshold; BVP, basic vegetative phase, the minimum duration of the vegetative phase above the threshold. Genotypes were sorted from short to long BVP.

| Genotype | Type | *ELF3A* | *GI* | *FTDel* | Group | DLc (h) | DLs (°Cd h<sup>-1</sup>) | BVP (°Cd) |
| --- | --- | --- | --- | --- | --- | --- | --- | --- |
| Sonali (Son) | Desi | elf3a* | GI96029* | Del | E | 10.9 | -182 | 722 |
| Genesis 509 (G509) | Desi | WT | GI96029* | WT* | D | 11.9 | -171 | 745 |
| PBA Slasher (Sla) | Desi | WT | GI96029* | WT* | D | 11.5 | -185 | 751 |
| PBA Striker (Str) | Desi | WT | GI96029* | WT* | D | 10.9 | -284 | 767 |
| PBA Maiden (Mdn) | Desi | WT | GI96029* | WT* | D | 10.9 | -276 | 780 |
| Neelam (Nee) | Desi | WT | GI96029* | WT* | D | 11.4 | -184 | 782 |
| PBA Monarch (Mon) | Kabuli | WT | WT | Del | A | 12 | -134 | 788 |
| PBA Royal (Roy) | Kabuli | WT | GI96029* | Del | C | 11.3 | -205 | 788 |
| CICA1841 (C841) | Desi | WT | GI96029* | WT* | D | 11.6 | -180 | 792 |
| Genesis 836 (G836) | Desi | WT | GI96029* | Del/WT*(a) | C/D | 10.7 | -437 | 794 |
| PBA Captain (Cap) | Desi | WT | GI96029* | Del | C | 11 | -239 | 801 |
| Howzat (Hwz) | Desi | WT | GI96029* | WT* | D | 11 | -280 | 804 |
| Kyabra (Kya) | Desi | WT | WT | Del | A | 11.4 | -208 | 816 |
| PBA Seamer (Smr) | Desi | WT | GI96029* | Del | C | 11.3 | -221 | 819 |
| Genesis 090 (G90) | Kabuli | WT | GI96029* | WT* | D | 10.8 | -416 | 820 |
| PBA Drummond (Dru) | Desi | WT | GI96029* | Del | C | 11.3 | -207 | 823 |
| PBA HatTrick (Hat) | Desi | WT | GI96029* | WT*(a) | D | 11.1 | -227 | 828 |
| CICA1812 (C812) | Desi | WT | GI96029* | Del | C | 10.7 | -436 | 828 |
| Jimbour (Jim) | Desi | WT | WT | Del | A | 11.2 | -235 | 834 |
| Tyson (Tys) | Desi | WT | GI96029* | Del | C | 11.5 | -153 | 835 |
| Amethyst (Amt) | Desi | WT | WT | Del | A | 11.5 | -153 | 837 |
| PBA Magnus (Mgn) | Kabuli | WT | WT | Del | A | 11.1 | -199 | 849 |
| Kaniva (Kan) | Kabuli | WT | WT | WT* | B | 11.4 | -174 | 850 |
| Yorker (Yrk) | Desi | WT | GI96029* | Del | C | 11.3 | -201 | 860 |

(a) Genotyping of the field material for PBA HatTrick and Genesis 836 at *FTDel* was different to the reference samples and published or publicly available information for these varieties (Del for both varieties; Nguyen et al., 2022; Home - Chickpea BLAST portal (adelaide.edu.au)). Assays on >50 individuals revealed that our field-phenotyped Genesis 836 was mixed at the *FTDel* locus

### Phenotyping

#### Phenology

Crops were monitored weekly or bi-weekly to determine the date when 50% of a plot reached: emergence (shoot visible above the soil level), flowering (at least one flower in the plant with separated petals), and podding (developing pods >2–4mm in length, surpassing the corolla). We used a thermal time scale for the duration of the main pheno-phases: from sowing to emergence, from emergence to flowering, and from flowering to podding. In all cases, thermal time was calculated from daily average temperature using a base temperature of 0 °C (Berger *et al*., 2006; Soltani *et al*., 2006). Physiological maturity was only scored in three environments.

#### Yield and its components

At harvest, we sampled a 0.5 m^2^ area in the central rows of the plots to assess shoot biomass, grain yield and its components at Kapunda and Merredin. We clipped all the plants at ground level within the sampled area and shoots were oven dried at 70 °C for 48 h. We weighed the dry samples before and after threshing to obtain shoot biomass, grain yield, and harvest index as the ratio between yield and biomass. We used a subsample of 400 grains to estimate the average grain weight, and from this value and grain yield, we estimated the number of grains per square meter. At Gatton, we only measured grain yield.

### Genotyping

The genetic background of the genotypes was characterised by their allelic composition for the relevant single nucleotide polymorphisms (SNPs) at the genes *CaELF3a* (hereafter *ELF3a*) and *CaGIGANTEA* (hereafter *GI*) (Ridge *et al*., 2017), and by the presence of a L30kb deletion on the *FTa1*-*FTa2* intergenic region described by Ortega *et al*. (2019) (hereafter *FTdel*) (Table 1). The allelic composition for *ELF3a* and *GI* was determined through High Resolution Melt using the primers given in Supplementary Table 1 and the following conditions: 2 μL of purified genomic DNA (50 ng μL^-1^) were used directly in 10 μL reaction volumes containing 5 μL of SensiFAST HRM mix (Bioline), 1.6 μL sterile distilled water and 0.7 μL of each primer (100 nM). PCR amplification, DNA melting and end point fluorescence level acquiring PCR amplifications were performed on a Rotor-Gene Q thermocycler (Qiagen, Hilden, Germany). PCR conditions as follows: 50 cycles of denaturation at 95 °C for 5 s, annealing at 60 °C for 10 s, extension at 72 °C for 20 s. High-resolution melting analysis was performed after PCR amplification with temperature ramping from 61 to 80 °C, rising by 0.1 °C 2s^-1^. *FTdel* was assessed with a PCR-based codominant maker. PCR was performed in a final volume of 25 μL containing 50 ng template DNA, 5 μL of 5x reaction buffer, 10 mM dNTPs, 0.2 μM of each primer (Supplementary Table 1), 50 mM MgCl_2_, 0.1 μL of MangoTaq^TM^ DNA polymerase (Bioline, Australia) and autoclaved Milli-Q water to final volume. Reactions were performed in a thermal cycler using the following program: an initial denaturation of 5 min at 94°C, followed by 40 cycles (94°C for 45s, annealing temperature of 58 °C for 30s, extension of 30 s at 72 °C) and a final extension of 10 min at 72 °C.

Genotyping was initially done on single, reference DNA samples of genotypes sourced from the Australian Grains Genebank (AGG). Field samples (n=3) were collected from the 2022 Kapunda field trial (early time of sowing), and their genotypes were compared with the reference material. Where genotypes did not match or were mixed for one or more genes, further sampling and genetic analysis was conducted to confidently determine allelic compositions. We note the importance of characterising actual material used for phenotyping, as seed of a given variety from different sources may sometimes genotype differently.

Genotypes were classified in five groups according to their allelic combination (Table 1): group A corresponds to all three late alleles; group B has only the early allele for *FTdel;* group C has only the early allele for *GI*; group D combines early alleles of *FTdel* and *GI*; and group E combines the early alleles of *ELF3a* and *GI*.

### Weather

Weather data including daily rainfall, maximum (Tmax) and minimum temperature (Tmin), relative humidity, vapour pressure deficit, and solar radiation were obtained from the nearest Australian Bureau of Meteorology station (www.data.longpaddock.qld.gov.au). To compare the growing conditions across environments, we considered a crop cycle spanning from sowing to 500 °Cd after onset of podding. Derived variables include daily mean temperature, as average of maximum and minimum; thermal amplitude, the difference between maximum and minimum temperature; the photothermal quotient, the ratio of solar radiation and mean temperature (Fischer, 1985); rainy days (rainfall ≥ 0.5 mm); days with frosts (Tmin ≤ 0 °C); and days with chilling (mean temperature ≤ 15 °C) or high temperatures (Tmax ≥ 35 °C). The threshold for chilling temperature is from Chauhan *et al*. (2022) and the threshold for high temperature is from Devasirvatham *et al*. (2012). Daylength, including civil twilights, was computed from sunrise and sunset timetables from the Astronomical Applications Department of the U.S. Naval Observatory (https://aa.usno.navy.mil/data/index).

### Data analyses

We used two-way analysis of variance (ANOVA) to assess the effects of the environment (E), genotype (G), and their interaction (G x E) on crop phenology and yield traits, where each environment resulted from the combination of site by season by sowing time. We computed the partial Eta squared (h2p) statistic to account for the size effect or the variance explained by E, G and G x E (Cohen, 2013). Although sowing date varied across environments, we ran additional ANOVA to qualitatively compare the effect of early versus late sowing for specific traits.

We quantified the genotype-dependent phenotypic plasticity of yield, biomass, and harvest index as the slope of the linear reaction norm relating the trait for each genotype and the environmental mean of the trait (Finlay and Wilkinson, 1963; Sadras and Richards, 2014).

Linear and non-linear models were used to probe for relations between the duration of the pheno-phases and environmental variables. For the phase from emergence to flowering, we used a 3-parameter segmented linear model to describe a typical daylength response of a long-day plant (Major, 1980). Parameters were DLc, the threshold daylength below which flowering time responds to daylength; DLs, daylength sensitivity, quantified as the rate of change in flowering time per unit change in daylength above the threshold; BVP, basic vegetative phase, the minimum duration of the vegetative phase above the threshold. Fig. 2c illustrates these parameters. As the duration of the juvenile phase (an early, daylength-insensitive stage) was unknown, we ran our model iterating daylengths from different times during the vegetative phase at 100 °Cd intervals. The best fit was obtained at 500 °Cd after emergence, so we used the daylength at this time in our model. Similarly, we correlated environmental variables for 100 °Cd windows along the flowering-to-pod set interval to identify the best descriptors for the duration of this phase across environments.

We used correlation, regression analysis, and correlation-based principal components analysis (PCA) to relate phenological and yield traits with the genetic background of the genotypes (*i.e.,* the allelic composition for *ELF3a*, *GI* and *FTdel*). Phenological traits included the mean thermal time duration of the phenological phases for each genotype, and the three parameters of their daylength response. Yield traits included the trait *per se* and their phenotypic plasticity.

## Results

### Weather conditions

Mean temperature during the growing season ranged from 11.7 °C in the early sowing of Kapunda 2021 to 17.1 °C in the late sowing of Gatton 2022. Kapunda was the site with lowest average temperature (12.3 ± 0.6 °C), followed by Merredin (13.3 ± 0.2 °C) and Gatton (15.8 ± 0.9 °C, Table 2). Average temperature was 0.8 ± 0.6 °C warmer in late-sown crops. Associated with these thermal conditions, the length of the growing season (from sowing to 500 °Cd after podding), varied 2-fold, from 101 days in the late sowing at Gatton 2021 to 201 days in the early sowing at Kapunda 2022 (Table 2). Chilling (frequency of days with mean daily temperature ≤15 °C) varied from 80 ± 6 % at Kapunda to 41 ± 10 % at Gatton. Frosts (Tmin ≤ 0 °C) were only registered in Merredin 2021 (7 events, 4 of which occurred in September during flowering of early sown crops) and in Kapunda 2022 (2 events in July, early in the vegetative phase). The number of days with Tmax > 35 °C was negligible in all environments.

**Table 2.** Weather conditions from sowing to 500 °Cd after podding in 12 environments resulting from the combination of sites, seasons, and sowing dates. Average (Tave), minimum (Tmin) and maximum temperature (Tmax) are daily means. Chilling and high temperature are the percentage of days with Tave ≤ 15 °C and Tmax ≥ 35 °C, respectively. Rain and reference evapotranspiration (ETo) are cumulative for the crop season, and fallow rain is the cumulative rain from 1 December of the previous year to sowing.

| Site | Season | Sowing | Growing<br>season (d) | Tave<br>(°C) | Tmin<br>(°C) | Tmax<br>(°C) | Chilling<br>temp.<br>(%) | High<br>temp<br>(%) | Rain<br>(mm) | ETo<br>(mm) | Fallow<br>rain (mm) |
| --- | --- | --- | --- | --- | --- | --- | --- | --- | --- | --- | --- |
| Gatton | 2021 | 1<sup>st</sup> | 123 | 15.2 | 7.6 | 22.8 | 49 | 0 | 122 | 328 | 457 |
|  |  | 2<sup>nd</sup> | 101 | 15.7 | 8.0 | 23.4 | 40 | 0 | 107 | 294 | 483 |
|  | 2022 | 1<sup>st</sup> | 122 | 15.1 | 8.1 | 22.2 | 49 | 0 | 270 | 318 | 1009 |
|  |  | 2<sup>nd</sup> | 116 | 17.1 | 10.2 | 24 | 26 | 0 | 295 | 377 | 1075 |
| Kapunda | 2021 | 1<sup>st</sup> | 176 | 11.7 | 6.4 | 17.1 | 87 | 0 | 350 | 341 | 67 |
|  |  | 2<sup>nd</sup> | 148 | 12.5 | 6.9 | 18.2 | 78 | 0 | 310 | 375 | 172 |
|  | 2022 | 1<sup>st</sup> | 201 | 12.1 | 7.5 | 16.7 | 83 | 0 | 509 | 391 | 81 |
|  |  | 2<sup>nd</sup> | 147 | 13.1 | 8.1 | 18.2 | 73 | 1 | 352 | 379 | 242 |
| Merredin | 2021 | 1<sup>st</sup> | 169 | 13.3 | 7.2 | 19.4 | 73 | 0 | 179 | 365 | 143 |

Rainfall during the growing season ranged from 107 mm in the late sowing at Gatton 2021 to 509 mm in the early sowing at Kapunda 2022. Compared to 2000-2022, actual seasonal rainfall was close to average at Kapunda and Merredin in 2021, and above average in the other environments. Rainfall in Kapunda was generally close to or higher than reference evapotranspiration (ETo). In Merredin, reference evapotranspiration doubled the amount of rainfall. Gatton has a summer-dominant rainfall, where winter-sown crops mostly rely on stored soil moisture. Both seasons in Gatton had rainy pre-sowing conditions, with 457 mm fallow rainfall in 2021, and 483 mm in 2022 (Table 2).

### Phenology

Thermal time from emergence to flowering, from flowering to podding, and from emergence to podding all varied in response to environment, genotype, and their interaction (Supplementary Tables S2-S4). The emergence-to-flowering phase varied 1.9-fold, from 611 °Cd to 1160 °Cd, with environmental effects accounting for 82% of the variance, and genotype and interaction each accounting for 7% of the variance (Fig. 1A, Supplementary Table S2). Emergence-to-flowering was shorter in late-sown crops in all environments.

**Figure 1:**
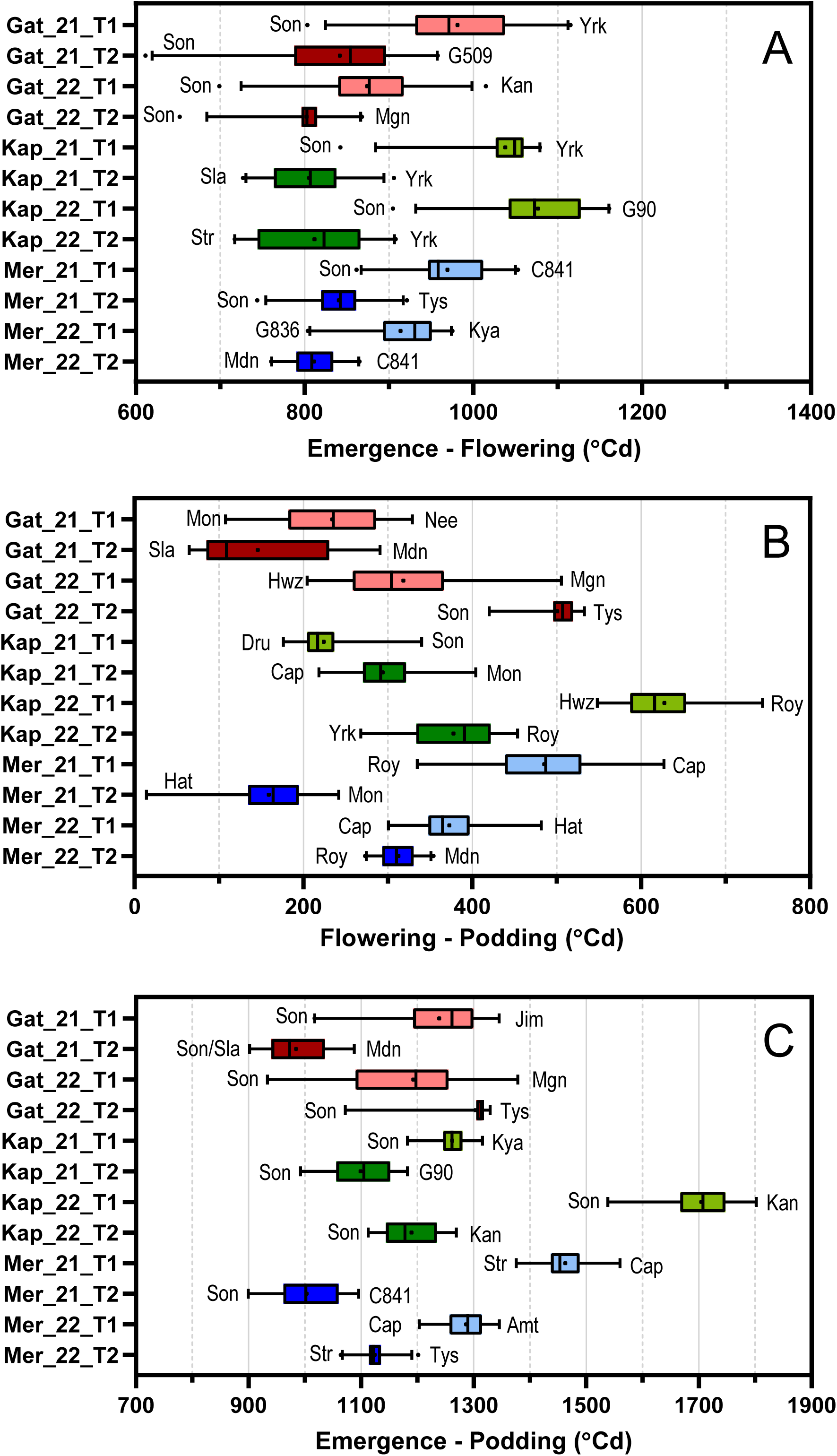
Thermal time from A) emergence to flowering, B) flowering to podding, and C) emergence to podding for 24 genotypes grown in 12 environments. Base temperature is 0 °C. Earliest and latest genotypes are shown next to the boxplots.

Variation in thermal time to flowering between the earliest and latest genotypes, ∼180 °Cd, was apparent in all the environments. Genotype rankings in flowering time changed across environments, consistent with the interaction (Fig. 1A; see also section “*Variation of thermal time from emergence to flowering…”*), but Sonali stood out as the earliest flowering genotype in 8 out of 12 environments, and Yorker was the latest to flower in 4 of the 12 environments.

The environment accounted for 89% of the variance in the flowering-to-podding interval, with small variation associated with genotype (3%) and interaction (4%) (Fig. 1B, Supplementary Table S3). This interval varied from 92 to 744 °Cd; it was shorter in late-sown crops at Gatton 2021 (100-200 °Cd) and longer in early-sown crops at Kapunda 2022 (> 600 °Cd). Variation with genotype in this trait was similar or higher than that of the emergence-to-flowering phase, with no consistent effect of time of sowing. None of the genotypes stood out for this trait.

The variation in the duration of both emergence-to-flowering and flowering-to-podding determined a substantial variation in the thermal time from emergence to podding, from 900 °Cd for late-sown Sonali at Gatton 2021 to 1802 °Cd in the early-sown Kaniva at Kapunda 2022 (Fig. 2C). Except for Gatton 2022, late sowing shortened the thermal time to podding. Sonali was the earliest genotype to set pods in Gatton and Kapunda and was in the top two earliest genotypes in 3 of the 4 environments of Merredin.

**Figure 2:**
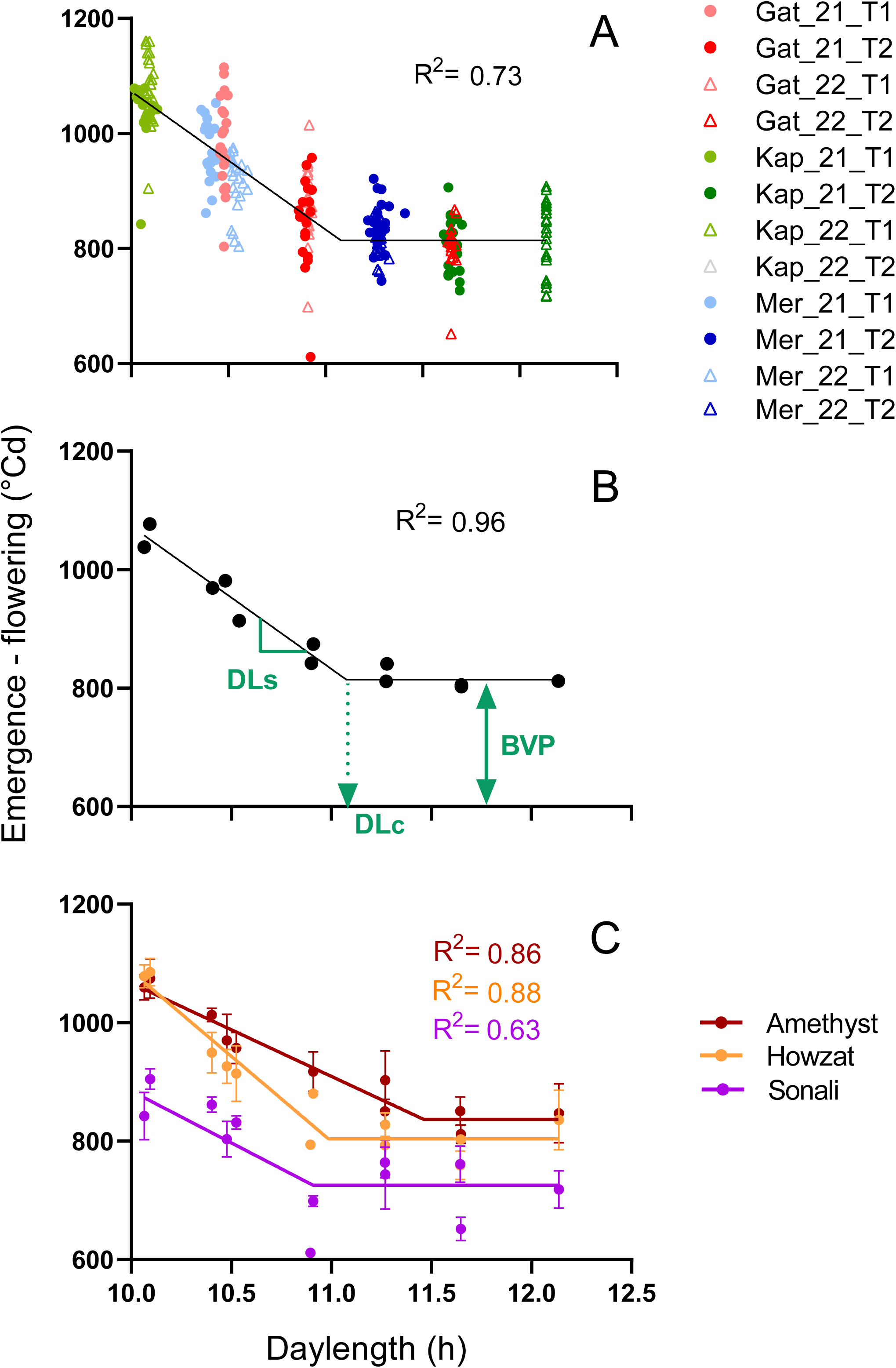
Association between thermal time from emergence to flowering and daylength in a model including A) all three sources of variation (environment, genotype, interaction); B) environment only; and C) a sample of three contrasting genotypes. The model has three parameters illustrated in (B): BVP: the minimum duration of the vegetative phase under inductive long-day conditions; DLc: the critical daylength below which flowering time delays increasingly with shortening daylength; and DLs: the daylength sensitivity (i.e, the delay in flowering time per hour of shortening daylength). The daylength is at 500 °Cd after crop emergence in each environment. In (C) error bars represent 1 standard error. All fitted models in A-C, had p < 0.05.

#### Genotype-dependent variation in flowering response to daylength

Across all three sources of variation, a 3-parameter, quantitative long-day response model accounted for 73% of the variation in thermal time from emergence to flowering (Fig. 2A). Averaged across genotypes, the model features: a critical daylength, DLc = 11.2 ± 0.1 h; a rate of change in flowering time below this threshold, DLs = 232.3 ± 17.4 °Cd h^-1^; and a minimum thermal time to flowering below the threshold, *i.e.,* basic vegetative phase, BVP = 804.7 ± 7.2 °Cd (Fig. 2B). All three parameters varied with genotype, causing changes in the rankings in flowering time across environments (Fig. 2C, Table 1 and Supplementary Fig. S1). For instance, Amethyst and Sonali had similar daylength sensitivity but differed in both critical daylength and basic vegetative phase. As a result, the difference in flowering time between them was larger in short-than in long-day. The difference in the three parameters of the daylength response between Amethyst and Howzat explained why Amethyst flowered slightly earlier than Howzat under short days (10 h), slightly later under long days (12 h), and much later under intermediate daylength (∼11 h) (Fig. 2B).

#### Variation in flowering-to-podding interval with environmental factors

The flowering-to-podding interval was unrelated to daylength (Fig. 3A). The interval lengthened with low radiation (Fig. 3B) and humid air conditions as captured in correlations with both relative humidity and vapour pressure deficit (Fig. 3C-D). Correlations with temperature-related variables were weaker or insignificant partially because of a few environments that departed from the general trend including Gatton 2022 and Kapunda 2021 early sowing (Fig. 3E-F, Supplementary Table S5, Supplementary Fig. S2). Excluding these environments, the flowering to podding interval correlated to both mean temperature (r = - 0.85, p<0.0001) and the frequency of chilling days after flowering (r = 0.74, p<0.0001). Correlations of the interval with other variables, like radiation or humidity, also improved after removing disruptive environments. As the disruptive environments were not the same for different variables, the combination of complementary variables would contribute to better explain the variation in the flowering-to-podding interval across environments. To test this hypothesis, we used the predicted interval from 3 regression models, each generated with one candidate variable (mean temperature, radiation and relative humidity) discarding their disruptive environments, and combined them in two approaches: a) a ‘balanced’ approach that averages the predicted interval from the three regressions to account for an integrated effect of the 3 variables, and b) a ‘strongest constraint’ approach, that kept only with the maximum estimated interval from the three regressions, to account for the variable with the highest effect on each situation (Fig. 4). Both approaches gave reasonable estimates for the interval across most of the environments, except for the early sowing of Kapunda 2021, which was disruptive for most of the variables.

**Figure 3:**
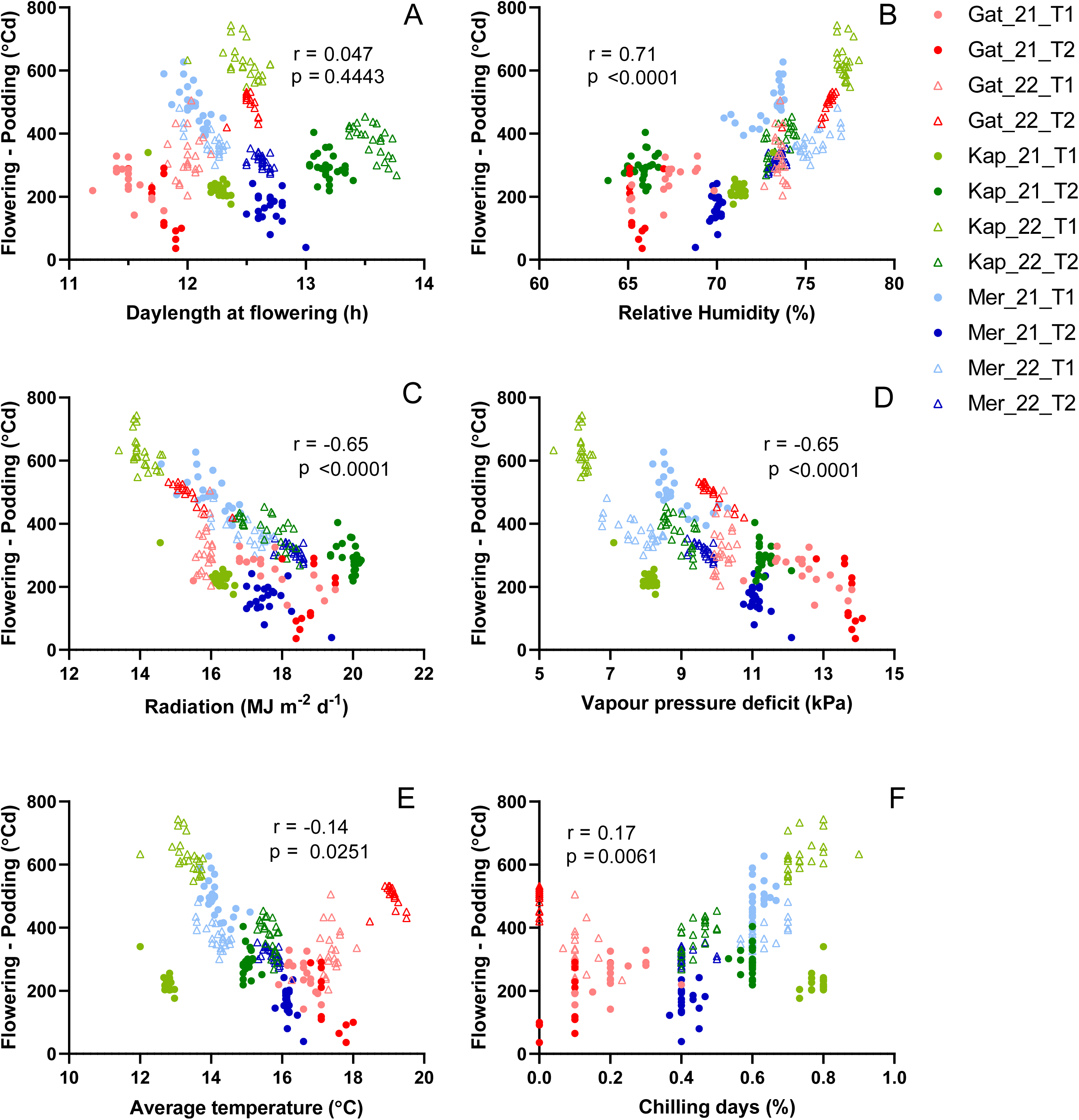
Relation between the flowering-to-podding interval and different environmental variables between flowering and 600 °Cd after flowering: A) Daylength, B) relative humidity, C) radiation, D) vapour pressure deficit, E) average temperature, and F) frequency of days with chilling temperature. Symbols are environments and the scatter for each environment is associated with 24 chickpea genotypes.

**Figure 4:**
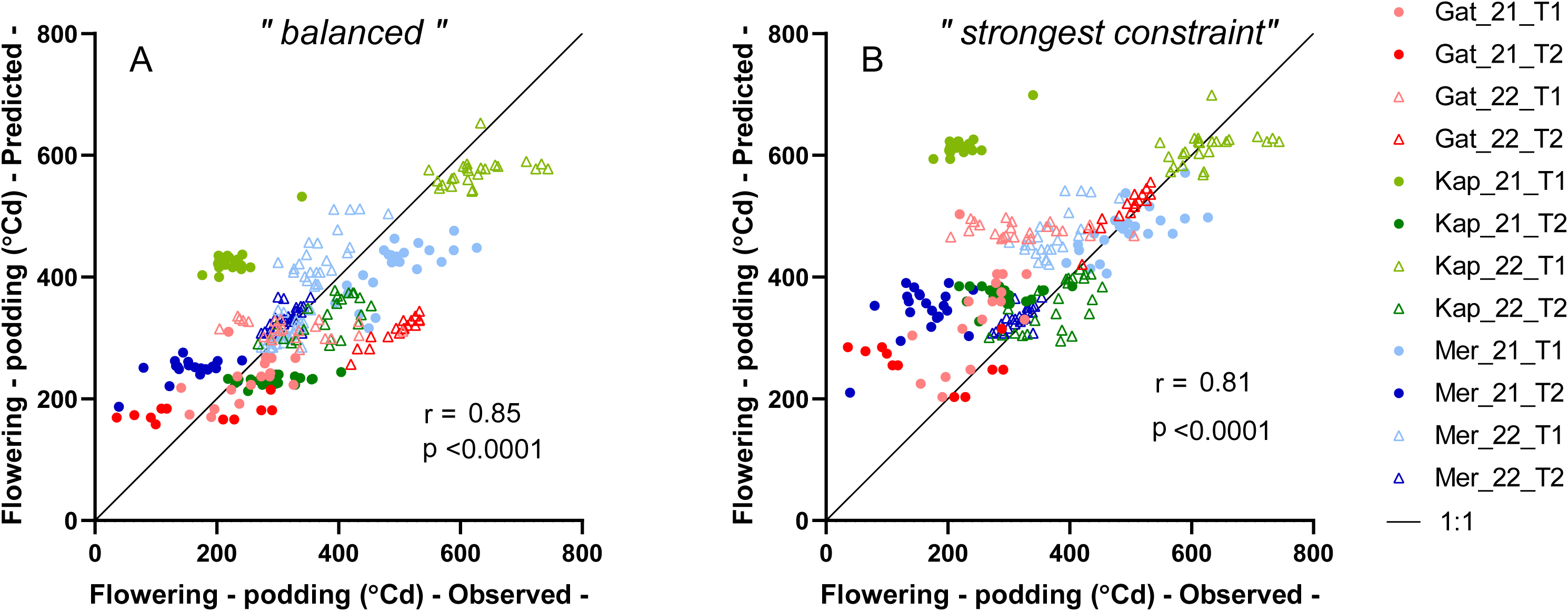
Observed vs modelled flowering-to-podding interval, using 2 approaches: A) the ‘balanced’ approach used the average predicted values from regressions of the interval vs three candidate constraints to pod set (mean temperature, radiation, and relative humidity); B) the “maximum constraint” approach, kept only the maximum value from the 3 regressions to account for the “most limiting” condition to pod set. Correlation coefficients are for 11 environments (Kap_21_T1 was not considered).

### Crop yield and its associations with biomass and harvest index

Grain yield ranged from 0.05 to 5.8 t ha^-1^ and varied with all three sources of variation (all p < 0.001; Supplementary Table S6). The environment accounted for 80% of the variance in yield, with 8% attributable to the interaction and 1% to genotype. Highest yields were recorded in Gatton 2021 (> 4 t ha^-1^), followed by Kapunda with intermediate and highly variable yield (2-4 t ha^-1^), and lower yield (< 2 t ha^-1^) at Merredin and Gatton 2022 (Fig.5A). Yield response to time of sowing was variable: late-sown crops either outyielded (p < 0.01, in Gatton 2021, Kapunda 2022 and Merredin 2021), under-yielded (Kapunda 2021 and Merredin 2022), or did not differ from early sown crops (p > 0.05, in Gatton 2022). No genotype was the best or worst yielding in more than two environments.

**Figure 5:**
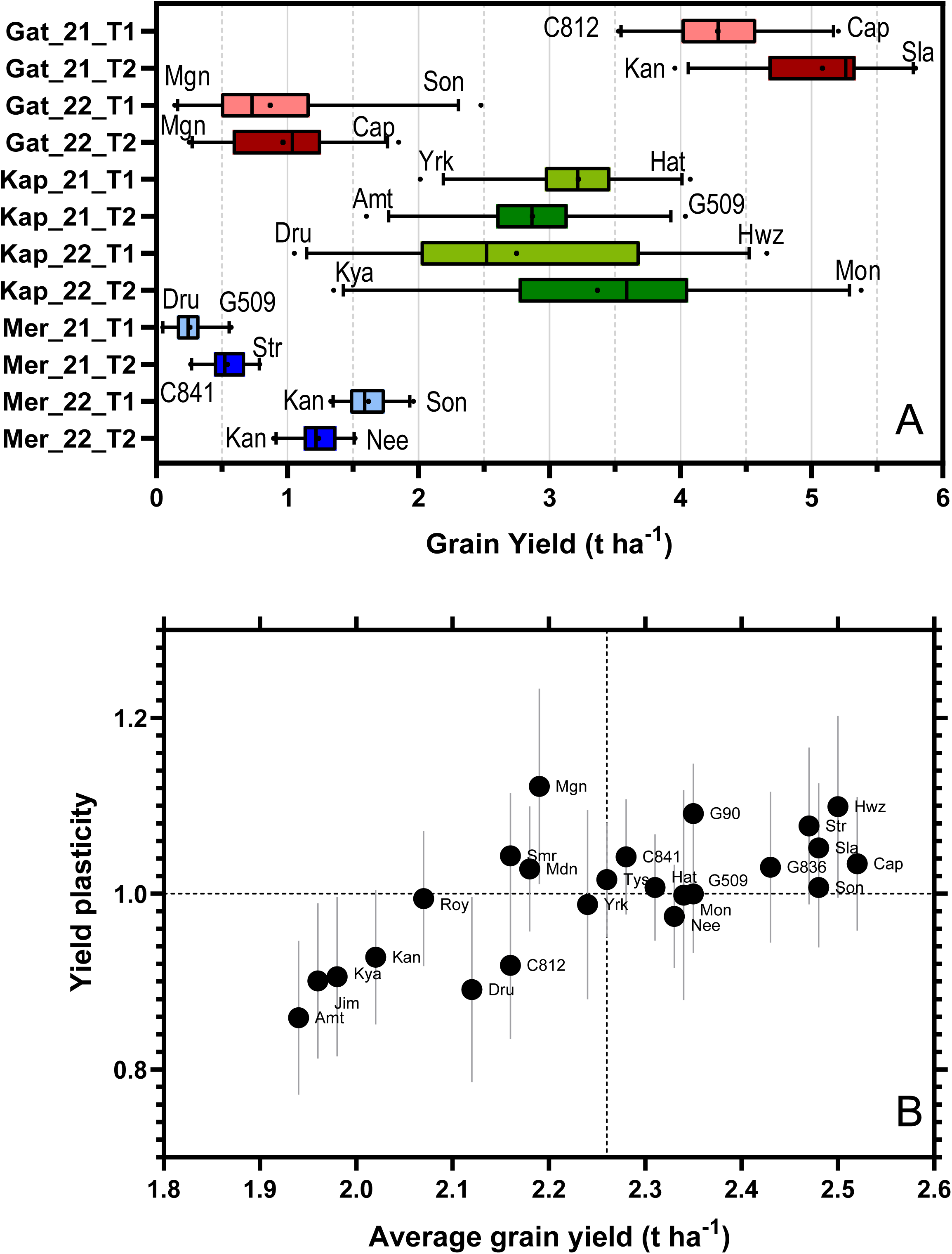
A) Grain yield of 24 chickpea genotypes across 12 environments; B) Phenotypic plasticity of yield as a function of average yield of each genotype across the environments. Labels show the lowest and highest yielding genotypes in A and all genotypes in B. In B, error bars are the standard error of the plasticity; the dashed lines separate four quadrants with combinations of higher- or lower-than-average yield (right and left quadrants) and higher- or lower-than-average plasticity (top and bottom quadrants).

Figure 5b depicts the alignment between the phenotypic plasticity of yield and the average yield across all the environments. The average yield (x-axis) varied from ∼2 t ha^-1^ for the lower yielding genotypes Amethyst, Jimbour, Kyabra and Kaniva, to ∼2.5 t ha^-1^ for the higher yielding Howzat and PBA Captain (p < 0.05, Supplementary Table S7). The plasticity (y-axis) ranged from 0.86 in Amethyst to 1.12 in PBA Magnus. Genotypes with high average yield and plasticity (PBA Captain, Howzat and Slasher in the upper-right quadrant) had superior yield in the best environments (Gatton 2021) while Sonali, with a high yield but an average plasticity, had good yield across the environmental gradient including the low-yielding environments like Gatton and Merredin 2022. Genotypes with low yield and low plasticity, like Amethyst, Jimbour, Kyabra and Kaniva (in the bottom-left quadrant), were the worst performing, especially in high yielding environments.

The analysis of shoot biomass and harvest index was limited to the Mediterranean-type environments of Merredin and Kapunda. Environment accounted for 89% of the variance of biomass, with a marginal effect of genotype and no interaction (Supplementary Table S8). Biomass correlated with rainfall during the crop cycle (Fig 6A), where the wet conditions for early-sown crops of Kapunda 2022 produced 15 times more biomass than the driest condition of late-sown crops at Merredin 2021 (Fig 6C). Early sowing produced, on average, 8.4 t ha^-1^ compared to 5.6 t ha^-1^ in their late-sown counterparts (p < 0.001). The large environmental variation in shoot biomass eclipsed genotype differences despite an average difference of more than 2 t ha^-1^ between Monarch (8.2 t ha^-1^) and Sonali (5.7 t ha^-1^, Supplementary Table S9). Furthermore, phenotypic plasticity of biomass varied with genotype (Fig. 6B), from ∼0.7 in Sonali, Maiden and Kyabra to 1.23 in Howzat (p < 0.05).

**Figure 6:**
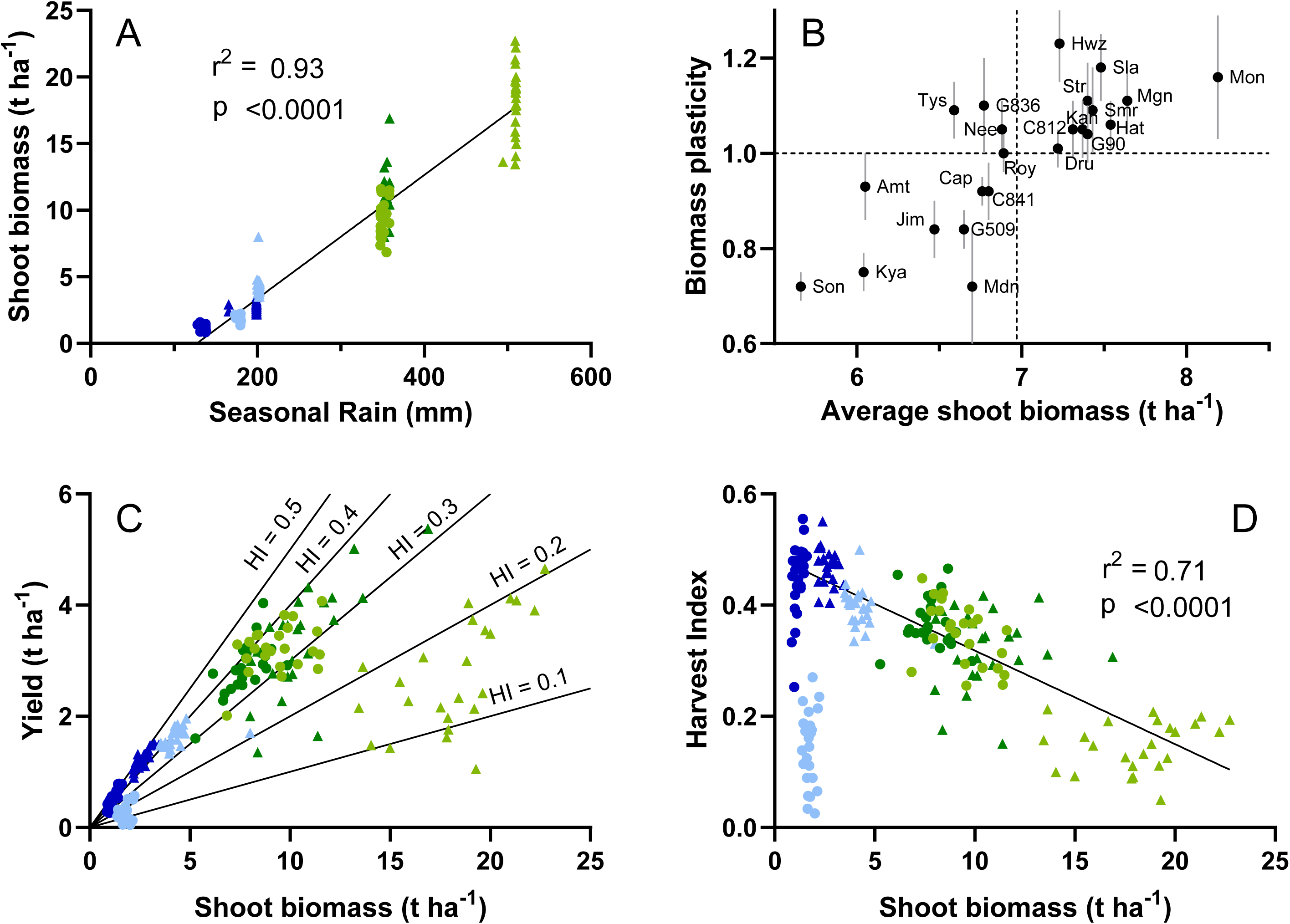
A) Association between shoot biomass and seasonal rainfall; B) Phenotypic plasticity (Y-axis) and average (X-axis) values of shoot biomass for each genotype across the environments (vertical segments indicate the standard error of the phenotypic plasticity); C) Association between grain yield and shoot biomass; and D) Association between harvest index and biomass. Data for 24 genotypes across 8 winter-rainfall environments, where the scatter is variation with genotype. Black lines in A and D, are linear regressions of the data (excluding Mer_21_T1 in D), and isolines of harvest index in (C).

Higher biomass did not directly translate into higher yield as harvest index varied widely with all three sources of variation (Fig. 6C and Supplementary Table S10). Across environments, harvest index varied from ∼0.14 in early-sown crops of Merredin 2021 or Kapunda 2022 to ∼0.45 in late-sown crops of Merredin in both seasons (Fig. 6C-D). In general, Kapunda crops had higher biomass than those of Merredin and late sowings had lower biomass but higher harvest index than early ones; harvest index increased from 0.26 for early sowing to 0.40 for late sowing (p < 0.001). Excluding the early sowing at Merredin 2021, where harvest index was likely affected by frosts during early reproductive stages (see section “Weather conditions”), harvest index correlated negatively with biomass (Fig. 6D). Among genotypes, harvest index ranged from an average of 0.40 in Sonali to 0.27 in Drummond (Supplementary Table S11).

### Associations between phenology and crop yield

Yield did not directly relate to any phenological trait across environments. Across sources of variation, harvest index declined with later podding (Fig. 7). Combining genotypes and environments, harvest index reduced 4.8% each 100 °Cd delay in podding from ∼0.5 when podding started 900 °Cd after emergence to ∼0.1 when it occurred at 1800 °Cd (Fig. 7A). This relationship persisted for environmental (Fig. 7B) and genotype-dependent variations, where early podding genotypes, like Sonali had higher harvest index than late-podding ones like Kaniva (Fig. 7C). The onset of pod set correlated to both the duration of the vegetative phase (r = 0.74, p < 0.001) and the flower-to-pod interval (r = 0.77, p < 0.001), which were modulated by different factors.

**Figure 7:**
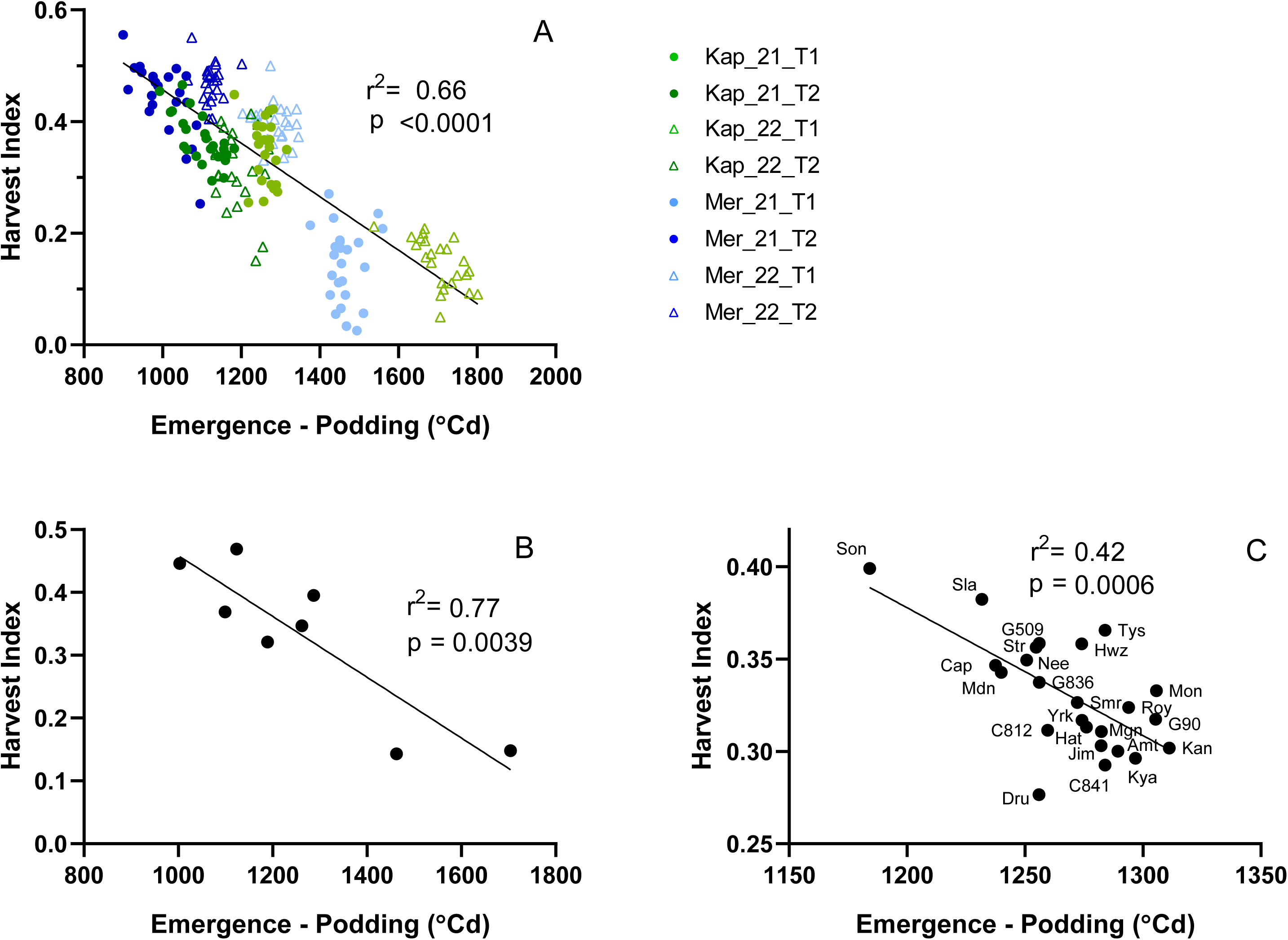
A) Association between harvest index and the time of podding in 24 chickpea genotypes grown in 8 environments. Data averaged by (B) environment and (C) genotype.

In a reduced dataset of the 3 environments where maturity date was recorded, both shoot biomass and yield correlated more to the length of the crop cycle, from emergence to maturity (r=0.92 and 0.88, for biomass and yield respectively) and to the duration of the pod setting phase (r=0.85 and 0.84, for biomass and yield respectively) than to time to flowering or podding (Supplementary Table S12). In this set of conditions, which did not include important variations in the flowering-pod interval or podding, grain yield strongly correlated to biomass (r=0.90, p < 0.001) and weakly with harvest index (r=-0.34, p<0.01).

### Associations between phenology and yield traits with genetic background

Principal component analysis integrated the phenological, growth, yield, and genetic data in winter-rainfall environments, capturing 54.6 % of the variance in two components (Figure 8). The first component (PC1, 31.1 % of the variance) correlated negatively with the thermal time to flowering (r = -0.92, p < 0.0001, Supplementary Table S12) and podding (r = -0.83, p < 0.0001), and positively with harvest index (r = 0.88, p < 0.0001) and yield (r = 0.72, p < 0.0001), reflecting the negative relationship between time of reproduction and harvest index (r=-0.75, p < 0.0001). The second component (PC2, 23.5 % of the variance) mainly correlated with shoot biomass (r = 0.88, p < 0.0001), and with both yield *per se* (r = 0.64, p < 0.01) and yield plasticity (r = 0.74, p<0.0001). Grain yield correlated weaklier with shoot biomass (r = 0.52, p < 0.01) than with harvest index (r=0.68, p < 0.001).

**Figure 8:**
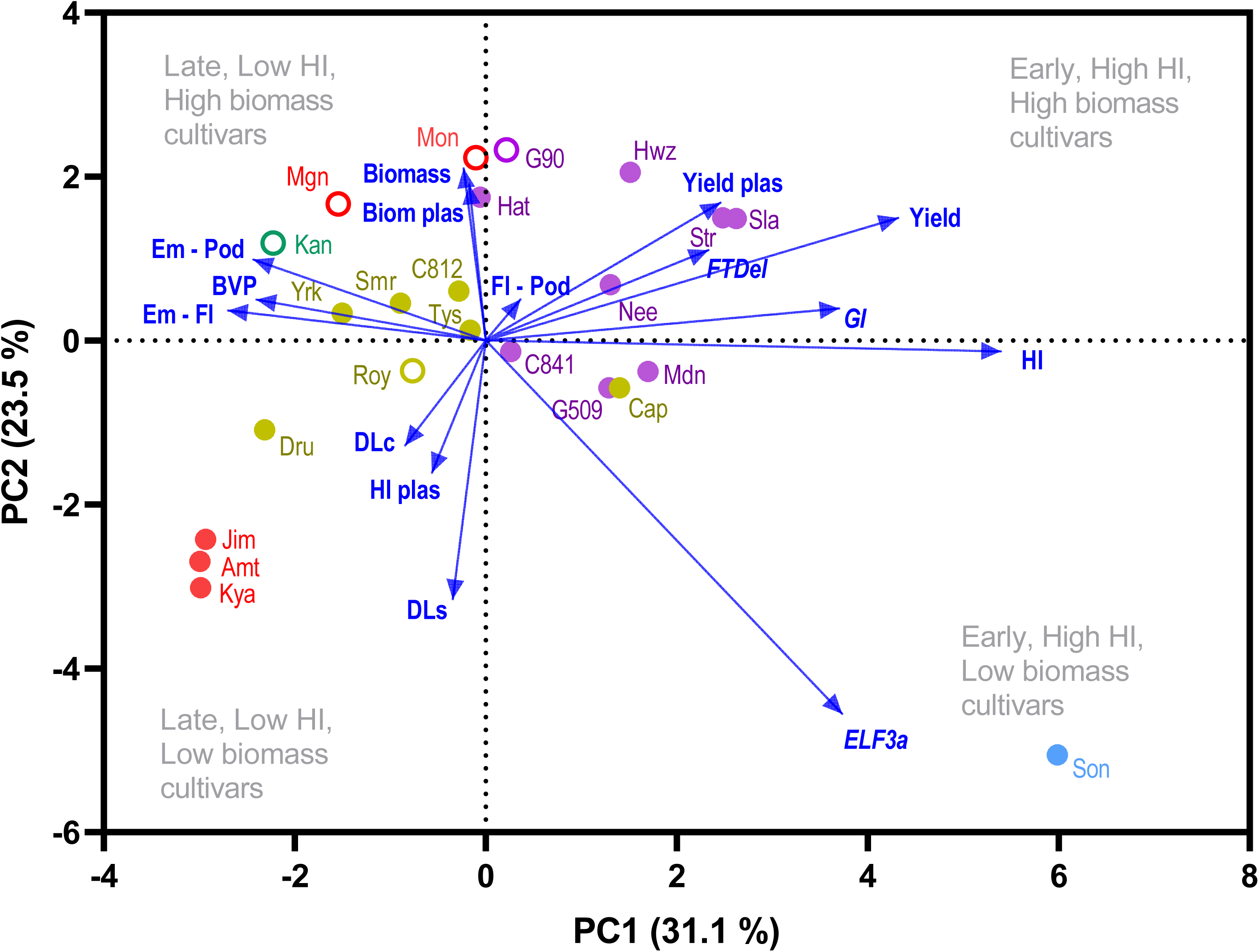
Principal components analysis of phenology and yield traits and the genetic background of 23 chickpea genotypes averaged across 8 environments. Vectors represent factor loadings in PC1 and PC2, and markers the PC scores for each genotype. Filled and open markers, represent desi and kabuli types, respectively. Traits are: Biomass (shoot biomass), Biom plas (plasticity of shoot biomass), Yield (grain yield), Yield plas (plasticty of grain yield), HI (harvest index), HI plas (plasticity of harvest index), Em - Fl (thermal time from emergence to flowering), Em - Pod (thermal time from emergence to podding), BVP (basic vegetative phase), DLs (daylength sensitivity), DLc (critical daylength) and the early allele of three candidate phenology loci: ELF3a, GI and FTDel. The genetic background of the genotypes, obtained from the allelic combination of the three loci, is shown with colours: group A: red, group B: green, group C: yellow, group D: violet, group E: light blue. Genesis 836 was not included in the analysis due to its mixed genotyping at the *FTDel* locus.

The biplot of PC1 and PC2 clustered early genotypes with high harvest index (right quadrants) and late genotypes with low harvest index (left quadrants) on the horizontal plane, and high and low biomass genotypes on the vertical plane (top and bottom quadrants, respectively). Sonali stood out as a small biomass, early-flowering and early-podding genotype with high harvest index (bottom-right quadrant). Kabuli genotypes concentrated in the opposite quadrant (top-left), with high biomass, late-flowering phenotypes, with low harvest index. High biomass, early flowering genotypes, that tended to be outstanding in good environments (*i.e.,* Howzat, Slasher and Striker), situated in the top-right quadrant, while the worst performing genotypes, characterised by late-flowering and small biomass (Kyabra, Amethyst, and Jimbour) situated in the opposite, bottom-left quadrant.

*ELF3a* showed high association with phenology and yield traits. The early allele of this gene was only present in Sonali, which clearly departed from the rest of the genotypes on both PC1 and PC2 dimensions. This allele correlated with early reproduction (r = -0.68 for thermal time to flowering and r = -0.77 for thermal time to podding, both p < 0.001), a short basic vegetative phase BVP (r = -0.50, p< 0.05), high harvest index (r = 0.49, p < 0.05) and low biomass (r = -0.49, p < 0.05). Individual associations of *GI* and *FTdel* with phenology and yield traits were weaker, but the combination of early alleles on both genes in Group D, correlated positively with PC1 (r = 0.54, p < 0.01) and grain yield (r = 0.47, p < 0.05), and negatively with BVP (r = -0.45, p< 0.05) and daylength sensitivity (r = -0.48, p< 0.05). The five genetic groups resulting from the allelic combinations, clustered in the PCA biplot: the three desi genotypes of Group A, that combines late alleles for the three loci, were late genotypes with low harvest index, biomass and yield. The two kabuli genotypes of Group A, PBA Monarch and PBA Magnus, had high biomass but intermediate to low harvest index, similar to Kaniva, the only genotype in Group B, that included only the early allele for *FTdel*. Most of Group C genotypes, that included only the early allele for *GI*, scored negatively for PC1 having intermediate to late phenology, intermediate to low harvest index and intermediate biomass. All Group D genotypes, that combined early alleles for both *GI* and *FTdel*, had positive scores for PC1, and combined early phenology and high harvest index with intermediate to high biomass. Group E, only represented by Sonali combining early alleles for *ELF3a* and *GI*, was the earliest genotype with the highest harvest index but the lowest shoot biomass.

## Discussion

High yield in grain crops requires the pairing of genotype and sowing date to avoid stressing factors in the critical period for grain set (Hunt *et al*., 2019; Lake and Sadras, 2014). This requires not only knowledge of the environmental constraints and their expected timing and intensity in each growing environment, but also on how genotypes modulate their phenology in response to the environment. This paper revisits these concepts for chickpea, in which phenology and particularly pod set have proven challenging for both breeders and agronomists.

### Environmental controls of flowering and flowering-to-podding interval

The manipulation of flowering time is crucial given that environmental conditions during the reproductive phase have a major impact on grain yield (Anwar *et al*., 2003; Nelson *et al*., 2010; Summerfield *et al*., 1996; Verghis *et al*., 1999). Besides temperature, which is implicit in the thermal time computation, daylength modulates flowering time in chickpea. In this work, we provide parameters for the daylength response of 24 chickpea genotypes, phenotyped under agronomically relevant conditions, which can be used to estimate flowering time (Ellis *et al*., 1994). These parameters can be readily incorporated into crop simulation models like APSIM, increasing the portfolio of available genotypes (Robertson *et al*., 2002b), or used to develop simpler, stand-alone applications to support farming decisions (*e.g.,* “Canola Flowering Calculator”, https://www.canolaflowering.com.au/). These parameters are physiologically meaningful traits that could be mapped to specific genomic regions (Pérez-Gianmarco *et al*., 2020) to provide a genetic basis for the differential flowering responses across environments (Lawn *et al*., 1995; Slafer and Rawson, 1996). Here we found, for example, partial correlations between the base vegetative phase BVP with both the early allele of *ELF3a* and the combination of early alleles of *GI* and the *FTa* cluster in the genetic Group D. As critical daylength DLc and daylength sensitivity DLs were correlated, but were unrelated to BVP, there may be scope to manipulate these traits separately to tailor phenology for specific environments (Slafer and Rawson, 1996).

The critical daylength was relatively low and showed little variation among genotypes, from 10.7 to 12.0 h, in comparison to other winter crops like wheat (DLc: 15-21h, Slafer and Rawson, 1996) or canola (DLc 15-17h, Robertson *et al*., 2002a). These DLc are similar to those used in different simulation models (Anwar *et al*., 2022; Chauhan *et al*., 2019; Singh and Virmani, 1996), but lower than DLc >15h reported in earlier experimental work for cultivated chickpea (Ellis *et al*., 1994; Roberts *et al*., 1985; Soltani *et al*., 2006) and wild relatives (Sharma and Upadhyaya, 2019). The practical meaning of a low DLc is that the impact of daylength regulating flowering time is restricted to early sowings and/or high latitudes, whereas its impact may be insignificant in low latitudes and/or late sowings (Vadez *et al*., 2013), as evidenced by the small differences in flowering time of the six late sowings across sites and years. This partly explains the undervalued role of daylength modulating flowering time in other studies (Li *et al*., 2022; Singh *et al*., 2021).

The flowering-to-podding interval varied widely across environments. Compared to other crops, where fruit set occurs shortly after flowering, this interval varied more than 60 days, from less than 100 °Cd to more than 800 °Cd in our study. Such long intervals were reported in environments of Australia (up to 30 days, Berger *et al*., 2004) and India (up to 50 days, Berger *et al*., 2006). Resources (water, nutrients, radiation, CO_2_) are the primary drivers of growth whereas non-resource factors including temperature, daylength and light quality are the main drivers of development (Connor *et al*., 2011). Pod setting has elements of both growth and development, but the variation in the flowering-to-podding interval mostly associates with environmental factors that cause the abortion of reproductive structures (Berger *et al*., 2004). Early-sown crops at Kapunda 2022 had 13.7 ± 1.8 aborted reproductive nodes below the first pod in comparison to 5.6 ± 1.0 in their late-sown counterparts. The environmental causes of abortion are partially understood (Singh *et al*., 2021), mainly attributed to chilling temperatures after flowering (Croser *et al*., 2003; Nayyar *et al*., 2005) and, less frequently, to other factors such as rainy days, overcast conditions, low light intensity, high soil moisture or high relative humidity (Croser *et al*., 2003; Rachaputi *et al*., 2021; Srinivasan *et al*., 1998; Verghis, 1996). As these weather factors are normally correlated in space and time (Bonada and Sadras, 2015; Sadras and Dreccer, 2015), crop response likely evolved integrating information from multiple environmental cues (Aphalo and Sadras, 2022). Our results indicate that multiple conditions may be required for pod set; for instance, crops at Gatton 2022 experienced non-chilling temperature after flowering, but low radiation and/or high relative humidity might have delayed pod setting. Conversely, the early-sown crops in Kapunda 2021 grew under high radiation and low humidity, but temperature <16 °C might have delayed podding (Berger *et al*., 2006). Understanding and integrating these variables into simulation or probabilistic models should contribute to a better prediction of the onset of pod set in chickpea, leading to better estimations of harvest index and grain yield (Aphalo and Sadras, 2022).

### Genetics of phenological development

The genetic basis of floral transition is best described in the model species *Arabidopsis thaliana*, where the information from both internal and environmental cues is combined into several molecular pathways that converge on the major integrator gene *FT*, which promotes the expression of floral identity genes that influence flower development (Kinoshita and Richter, 2020). Among these, the photoperiodic flowering pathway is responsible for floral induction in response to daylength. Briefly, inputs from light receptors are integrated by the plant circadian clock network and signalled downstream through clock-associated genes like *GIGANTEA* (*GI*), which induce *FT* expression (Mizoguchi *et al*., 2005; Sawa and Kay, 2011; Takagi *et al*., 2023). This pathway is well conserved across the plant kingdom, including legumes (Lin *et al*., 2021; Maple *et al*., 2024; Weller and Macknight, 2018), and presumably chickpea. For example, the flowering gene *CaELF3a* is part of the evening complex, an integral part of the circadian clock that represses flowering under non-inductive daylength (Liu *et al*., 2001; Ridge *et al*., 2017). A non-functional *caelf3a* allele for this gene has been associated with early flowering under short days in cultivated chickpea (Ridge *et al*., 2017), similarly to *elf3* mutants in other legumes such as lentil, pea and soybean (Fang *et al*., 2021; Lu *et al*., 2017; Weller *et al*., 2012; Yue *et al*., 2017). Sonali, the only accession carrying the *caelf3a* allele in our study, showed a conspicuous early phenology across all environments. Interestingly, its earliness was attributable to a shorter basic vegetative phase rather than a lower daylength sensitivity. Since *elf3* mutants have been traditionally categorised as photoperiod insensitive, this observation offers a new insight about the inhibitory nature of this gene.

The role of *FT* orthologues as central hubs for signal integration remains a conserved feature in the legume flowering pathway. Among these, *FTa1* plays a pivotal role integrating photoperiod and vernalisation signals (Hecht *et al*., 2011; Laurie *et al*., 2011; Yuan *et al*., 2021; Yue *et al*., 2017). Notably, in chickpea, a substantial deletion (≈30Kb) in the intergenic region between *CaFTa1* and *CaFTa2* is proposed as the causal polymorphism responsible for inter and intra-specific phenological differences (Ortega *et al*., 2019). Here, the wild-type allele (without deletion) was associated with earliness. While this contrasts with the reported effect of the deletion in Ortega *et al*. (2019), it supports the deletion difference observed between cultivated varieties Rupali and Genesis 836 (Nguyen *et al*., 2022) and its association with the phenotype in a derivative population (Atieno *et al*., 2021). The extremely complex regulation of *FT* orthologues and their interactions with downstream proteins involved in the onset of flowering makes it challenging to predict the effect of sequence variation in regulatory regions, which might vary with the environment and genetic dosage at other loci influencing *FT* induction.

*GIGANTEA*, the third flowering gene tested in the present study, appears to have a consistent function linking the circadian clock with *FT* induction in legumes (Hecht *et al*., 2007; Watanabe *et al*., 2011). Beyond flowering, *GIGANTEA* is associated with adaptation to drought, cold and salinity, and with the accumulation of starch and chlorophyll (Mishra and Panigrahi, 2015). The pleiotropic nature of *GIGANTEA* may explain its association with both early phenology and higher biomass (Fig. 9) and opens the possibility for the simultaneous improvement of multiple traits. Such strategy has been successfully applied in soybean for the coupled selection of maturity and salt tolerance (Dong *et al*., 2022).

Many studies have identified four major loci and dozens of QTL contributing to the control of phenology in chickpea, so attempting to account for all the variation with the three candidate genes examined in this research would be overly simplistic. Despite its limitation, our findings under agronomically relevant conditions highlight their potential to supplement conventional breeding to manipulate chickpea phenology.

### Implications for breeding

Early reproduction has long been recognised as a desirable trait to achieve higher or more stable yields in chickpea in a range of environments (Berger *et al*., 2004; Mallikarjuna *et al*., 2019; Nunavath *et al*., 2020). Chickpea grain yield is closely related to grain number, which is in turn proportional to the duration and growth rate during this critical period (Lake and Sadras, 2016; Sadras *et al*., 2015). Where terminal drought constrains the duration of the critical period of pod set, early reproduction could lengthen the pod setting phase (Berger, 2007; Or *et al*., 1999; Soltani *et al*., 2005; Subbarao *et al*., 1995).

Although “early varieties” can be associated to different traits (*i.e.* early flower initiation, early flowering, early pod initiation, early maturity), the breeding target is usually early flowering (Anbessa *et al*., 2007; Kumar and Abbo, 2001; Mallikarjuna *et al*., 2019; Nelson *et al*., 2010; Nunavath *et al*., 2020). This is valid for most grain crops (Gaur *et al*., 2008), where the time of flowering and fruit setting are tightly correlated, but it may not be suitable for chickpea, where: i) the onset of pod setting is more correlated to grain yield traits such as harvest index than flowering time; ii) the flowering-to-podding interval highly variable; and iii) the environmental control for flowering time and pod setting are different (Fig. 2 vs Fig 3).

Selecting for early flowering alone would be of little value for environments where prevailing aborting factors may delay podding (Jettner *et al*., 1999). This is especially relevant for Mediterranean environments in Australia and southern Indian environments where chilling tolerance has been identified as a breeding priority (Berger, 2007) and where genotypes with improved tolerance have been released (Clarke *et al*., 2004; Rani *et al*., 2020). In our work Sonali, selected for apparent chilling tolerance, stood out as an early-flowering and early-podding phenotype in most of the environments. Although Sonali set pods earlier in the season (hence, under abortion-prone conditions), its flowering-to-pod interval did not differ from the rest, as observed elsewhere with other chilling-tolerant genotypes (Berger *et al*., 2004). This suggests that either the chilling tolerance is relatively limited in these genotypes (Berger *et al*., 2005; Berger *et al*., 2012), or that other environmental factors may also cause pod abortion as discussed in the previous section. The benefits of combining early flowering and podding were evident in India where crossbred lines combining both traits showed an improved harvest index (∼52%) compared to late flowering lines that avoided chilling exposure (∼40%) (Rani *et al*., 2020).

Delayed maturity may potentially extend the pod setting phase. Besides an enhanced chilling tolerance, Sonali and other genotypes were selected for early maturity to avoid terminal drought (Gaur *et al*., 2008). While this may be an advantage in drought-prone environments (Kaloki *et al*., 2019) there may be a yield trade-off in more favourable seasons and locations. In our study, Sonali combined high yield and average phenotypic plasticity of yield: it returned relatively high yield in water-limited environments at the expense of lower biomass and yield in wetter and longer seasons. Although it is likely that early podding and early maturity are linked, *i.e.*, pod production hastens crop maturity (Anbessa *et al*., 2007), the possibility of combining early podding with more indeterminate growth habit may be adaptive for environments like the southern high rainfall region, with cooler and wetter spring and early summer, where water deficit is a lesser limitation (Rao *et al*., 2023).

### Implications for crop management

Improvement of chickpea grain yield could be linked to an increase in biomass, an increased harvest index or both (Anwar *et al*., 2003). The relative importance of biomass and harvest index varies with the environment, and there is normally a trade-off between them (Berger *et al*., 2004). Phenology modulates this trade-off where early podding favours a higher harvest index at the expense of shoot biomass and vice versa (Anbessa *et al*., 2007). Our results on how environmental and genetic factors control the onset of pod setting help to interpret, and potentially manage this trade-off.

Understanding abortion factors and restricting conditions at the end of cycle (*e.g.,* terminal drought or extreme temperatures) is important. Early-sown crops normally grow under less inductive conditions (long days), have a longer vegetative phase, and present a lower harvest index than late-sown crops (Richards *et al*., 2022). This pattern is exacerbated in environments prone to flower abortion, where the delay in pod setting translates into an even lower harvest index. In this kind of environment, both delaying sowing to avoid abortion factors near flowering, and using genotypes with improved tolerance to chilling temperature (like Sonali), are strategies to enhance the partitioning of biomass to grain (Berger *et al*., 2004; Kaloki *et al*., 2019; Saxena and Johansen, 1988). Sowing too late, however, may result in heavy yield penalties, if the higher harvest index does not fully compensate for the lower shoot biomass. In addition, late-sown crops might have a less developed root system with potential implications for terminal drought. In less restrictive environments, where abortion factors do not constrain harvest index, earlier sowing and long-season genotypes like Howzat would make a better use of the seasonal resources to produce a higher biomass that translates into higher yield (Nelson *et al*., 2010; Saxena and Johansen, 1988).

### Implications for the definition of the critical period for grain yield

The notion of a species-specific critical period for seed production is central to both fitness in nature (Darwin, 1859) and crop production in agriculture (Li *et al*., 2022). Growth conditions during this developmental period modulate grain number, which is the main source of variation in yield (Sadras, 2021). Farmers combine genotypes and sowing time to manipulate crop phenology to avoid biotic and abiotic stress. So, a well-defined critical period and a proper characterisation of the growth conditions for grain set are both necessary for crop management. Our results suggest that caution is needed to characterise both elements in chickpea.

Using sequential shading, Lake and Sadras (2014) identified the critical period for yield determination in chickpea as a window spanning from 300 °Cd before to 500 °Cd after flowering. This definition has been useful to explain yield variation in field experiments (Lake *et al*., 2016; Lake and Sadras, 2016; Sadras and Dreccer, 2015), but may not be suited for environments prone to pod abortion, where the flower-to-pod interval is likely to extend. For instance, if we strictly follow this definition, the whole critical period for early sown crops in Kapunda 2022 would have occurred before the first pod set in the earliest genotype, i.e., 548 °Cd after flowering for Howzat. An alternative definition of the critical period, that refers to the onset of pod set instead of flowering time, would include environments prone to the abortion of reproductive organs, without affecting its utility nor its physiological meaning in other environments (Carrera; *et al*., 2023). This adjustment would make sense as not only yield traits, chiefly harvest index, correlated more strongly with podding time than with flowering time across environments, but also as it is the most sensitive stage within the critical period (Lake and Sadras, 2014). Based on Lake and Sadras (2014), and assuming a flower-pod interval of ∼150 °Cd for no abortion conditions (Chauhan *et al*., 2023), the critical period could be redefined as an 800 °Cd window bracketing the onset of podding.

The photothermal quotient, calculated as the ratio between radiation intercepted by the canopy and mean temperature, is a useful approach to quantify spatial-temporal differences in growth conditions during the critical period (Cantagallo *et al*., 1997; Magrin *et al*., 1993; Rivelli *et al*., 2023; Sadras and Dreccer, 2015). It correlates with grain number and yield because it captures the correlation between growth and photosynthesis associated with radiation, and the negative association between the duration of critical period and temperature (Cantagallo *et al*., 1997; Fischer, 1985; Magrin *et al*., 1993). In chickpea, however, the correlation between yield and photothermal quotient would break down where conditions such as chilling temperature compromises pod setting. Poggio *et al*. (2005) resolved this limitation by assuming a photothermal quotient = 0 for field peas under chilling temperature (mean temperature < 10 °C) during the critical period. A similar approach could be tested for chickpea considering not only chilling temperature, but also other environmental variables associated to pod abortion.

### Conclusion

Under realistic field conditions and with a sample of germplasm representing the current availability of cultivars to farmers, we showed the biological and agronomic significance of the onset of podding to complement the most common marker - time of flowering. We demonstrated the discrepancy in the factors modulating flowering and podding time and provided elements to improve the prediction of these phenological stages based on genotype-specific daylength responses for flowering, and a combination of factors causing flower abortion and delayed pod setting, with implications on harvest index. These insights can inform simulation models or stand-alone applications to predict phenology and can be combined with remote sensing quantification of crop growth to provide grain yield estimations. The associations between *CaELF3a*, the *FT* cluster on chromosome 3 and *GIGANTEA*, with field phenotypes of distinct phenology, shoot biomass and harvest index characteristics, supports the relevance of these loci for breeding.

## Supplementary data

The following supplementary data are available at JXB online.

*Supplementary Table S1. primers used for allelic discrimination*.

*Supplementary Table S2. ANOVA for the thermal time from emergence to flowering*

*Supplementary Table S3. ANOVA for the thermal time from flowering to podding*.

*Supplementary Table S4. ANOVA for the thermal time from emergence to podding*.

*Supplementary Table S5. Correlation matrix for the flowering-to-podding interval and 13 weather variables in alternative periods*.

*Supplementary Table S6: ANOVA for grain yield*.

*Supplementary Table S7: Average grain yield and its phenotypic plasticity for 24 chickpea genotypes*.

*Supplementary Table S8: ANOVA for shoot biomass*

*Supplementary Table S9: Average shoot biomass and its phenotypic plasticity for 24 chickpea genotypes*.

*Supplementary Table S10: ANOVA for the harvest index*.

*Supplementary Table S11: Average harvest index and its phenotypic plasticity for 24 chickpea genotypes*.

*Supplementary Table S12: Correlation matrix of yield and phenological traits for a subset of data including 3 environments with reliable maturity scorings*.

*Supplementary Table S13: Correlation matrix of yield and phenological traits, and the allelic composition of 24 genotypes averaged across 8 environments*.

*Supplementary Fig. 1: Flowering time as a function of daylength 500°Cd after emergence for 24 chickpea genotypes*.

*Supplementary Fig. 2: Correlation matrix for the Flower-Pod interval and 13 weather variables averaged from flowering to 600 °Cd after flowering*

## Acknowledgements

We thank the Grains Research and Development Corporation (GRDC) for investments supporting our fieldwork (UOT1909-002RTX) and RG’s PhD Scholarship; Steffan Schmitt (Agricultural Consulting and Research) and staff of DPIRD Merredin Research Facility for the sowing and maintenance of the crops; Tan Dang, Habtamu Tura, Han Chow and Ian Bogisch for field and laboratory work; Jason Brand and Arun Shunmugan (Agriculture Victoria) for seed and associated information; and Jakob Buttler for useful comments on the manuscript.

## Author contributions

RG, LL and VOS: conceptualization; RG, LL, MCC and VOS: method; RG, ROM, JH and VS formal analysis; RG, LL, ROM, JH, FD and BF: investigation; LL and MCC: resources; RG and LL: data curation; RG, ROM, JH and VOS: writing – original draft; RG, LL, MCC, ROM, JH, FD, BF, JW and VOS: writing – review & editing; RG and VOS: visualisation; LL, JW and VOS: supervision; JW and VS: funding

## Data availability

The data supporting the conclusions of this article will be made available by the authors, upon reasonable request.

